# Disparate pathways for extrachromosomal DNA biogenesis and genomic DNA repair

**DOI:** 10.1101/2023.10.22.563489

**Authors:** John C. Rose, Ivy Tsz-Lo Wong, Bence Daniel, Matthew G. Jones, Kathryn E. Yost, King L. Hung, Ellis J. Curtis, Paul S. Mischel, Howard Y. Chang

## Abstract

Oncogene amplification on extrachromosomal DNA (ecDNA) is a pervasive driver event in cancer, yet our understanding of how ecDNA forms is limited. Here, we couple a CRISPR-based method for induction of ecDNA with extensive characterization of newly formed ecDNA to examine ecDNA biogenesis. We find that DNA circularization is efficient, irrespective of 3D genome context, with formation of a 1 Mb and 1.8 Mb ecDNA both reaching 15%. We show non-homologous end joining and microhomology mediated end joining both contribute to ecDNA formation, while inhibition of DNA-PKcs and ATM have opposing impacts on ecDNA formation. EcDNA and the corresponding chromosomal excision scar form at significantly different rates and respond differently to DNA-PKcs and ATM inhibition. Taken together, our results support a model of ecDNA formation in which double strand break ends dissociate from their legitimate ligation partners prior to joining of illegitimate ends to form the ecDNA and excision scar.

**SIGNIFICANCE:** Our study harnesses a CRISPR-based method to examine ecDNA biogenesis, uncovering efficient circularization between DSBs. ecDNAs and their corresponding chromosomal scars can form via NHEJ or MMEJ, but the ecDNA and scar formation processes are distinct. Based on our findings, we establish a mechanistic model of excisional ecDNA formation.

## INTRODUCTION

Extrachromosomal DNA (ecDNA) drives accelerated tumor evolution and is associated with worse outcomes for patients with a wide variety of cancers (1–4). Potent oncogenes are frequently encoded on ecDNA and can become massively amplified in copy number and upregulated in gene expression(2,4). ecDNA is circular and acentric, ranging in size from 100 kb to several Mb(2,5). Due to their lack of centromeres, ecDNA are randomly segregated at cell division and drive intratumoral heterogeneity, facilitating rapid genomic changes to adapt to treatment or metabolic stress(6). Despite these advances, the biogenesis of ecDNA remains poorly understood.

Multiple mechanisms of ecDNA formation have been proposed, including circularization of DNA segments generated by two or more breaks on the same chromosome arm, which we term “excisional ecDNA formation”, or ligation of multiple segments resulting from chromothripsis. Mechanistic investigations of ecDNA formation have proved challenging, as current studies rely on patient samples, patient-derived cell lines, or cell lines subjected to drug selections (7–10). In these studies, the requirement for selection—either within the tumor microenvironment or *in vitro*—of rare cells possessing spontaneously-generated ecDNA obscures the process of ecDNA formation. More recently, ecDNA have been engineered in human cells and mice using Cre-lox, enabling examination of ecDNA without extended selection for ecDNA+ cells(11). However, due to the use of Cre-mediated recombination to form ecDNA, this method cannot be used to examine the process of ecDNA formation itself.

Mechanistic understanding of excisional ecDNA biogenesis has thus remained elusive. We consider two primary models for how excisional ecDNA formation may occur based on the literature: (1) the end-swapping model, and (2) the dissociation model. In the end-swapping model, the two double-strand breaks (DSBs) undergo synapsis, holding each end in close proximity to its legitimate partner. Subsequently, these two synapses come into close physical proximity, the repair foci merge, and the ends are swapped and ligated, generating the ecDNA and chromosomal excision scar. The end-swapping model is analogous to the mechanism of reciprocal translocations(12). In the dissociation model, a failure to form or maintain the DSB synapses is followed by dissociation of the legitimate DSB end partners away from one another. Approximation and ligation of the ecDNA and excision scar thus proceed as relatively independent processes. These two models yield divergent predictions for (1) the impact of chromosome conformation on circularization efficiency, (2) the relative rates of ecDNA vs. scar formation, (3) the similarity of repair signatures on ecDNA and scar junctions, and (4) whether inhibition of DNA damage response (DDR) proteins would differentially impact ecDNA vs scar formation. To test these predictions and distinguish between these models, there is a need for model systems that enable us to study ecDNA molecules immediately after formation. Further, such models would provide an opportunity to directly identify factors that may enhance ecDNA formation.

We recently applied CRISPR-C, a CRISPR-based method for generating circular DNA(13), to the study of early ecDNA evolutionary dynamics, quantifying the relationship between selection pressure strength and ecDNA abundance(6). CRISPR-C offers temporal control and flexibility in loci selection, while also relying on endogenous DNA repair pathways for DNA circularization. Here, we use CRISPR-C to examine the process of ecDNA biogenesis itself. Using droplet digital PCR (ddPCR), fluorescence microscopy, chromosome conformation capture (Hi-C), and targeted deep sequencing of ecDNA and scar junctions, we directly examine the early stages of ecDNA formation following excision from chromosomes. We find that circularization of megabase scale DNA fragments between DSBs is efficient. We show that ligation of the ecDNA and its corresponding chromosomal excision scar can be mediated by non-homologous end joining (NHEJ) or microhomology-mediated end joining (MMEJ). Pharmacologic inhibition of DNA-PK favors ecDNA formation, whereas ATM inhibition limits it. Collectively, these results validate the dissociation model of excisional ecDNA formation, shedding light on some of the molecular processes involved in its formation.

## RESULTS

### Generation of ecDNA with CRISPR-C

CRISPR-Cas9 can be delivered into cells to induce DNA breaks. Two breaks on the same chromosome can be ligated to form a DNA circle and leave behind an excision scar, a method termed CRISPR-C (**Fig 1A**). To validate the utility of CRISPR-C for the study of ecDNA, we investigated whether we could generate ecDNA containing MYC, one of the oncogenes most frequently amplified on ecDNA in human cancers(4). We designed CRISPR guides to excise a region that encompassed the entire chromosome 8 span of a MYC-containing ecDNA found in COLO320-DM(2), with two guides targeting the centromeric end and three guides targeting the telomeric end (**Fig 1A-C**). We delivered Cas9 with all pairwise permutations of centromeric and telomeric guide pairs into the near-haploid Hap1 cell line. Circularization was detected for all 6 guide combinations by inverse PCR (**Fig S1**). Quantitative analysis by ddPCR revealed mean circularization frequencies ranging from 1.12 to 2.89% (**Fig 1D**), while no circle junctions were detected when only a single centromeric guide was delivered. We repeated this experiment with 4 different pairs of guides to generate a 1.8 Mb DHFR-containing ecDNA, again detecting ecDNA for all guide pairs (**Fig 1E-F**).To confirm that CRISPR-C resulted in ecDNA rather than other structural rearrangements, we performed DNA fluorescence in-situ hybridization (FISH) on metaphase chromosome spreads and validated the extrachromosomal localization of CRISPR-C excised ecDNA (using DHFR L1 and R2 sgRNAs(6)) (**Fig 1G**).

**Figure 1.**
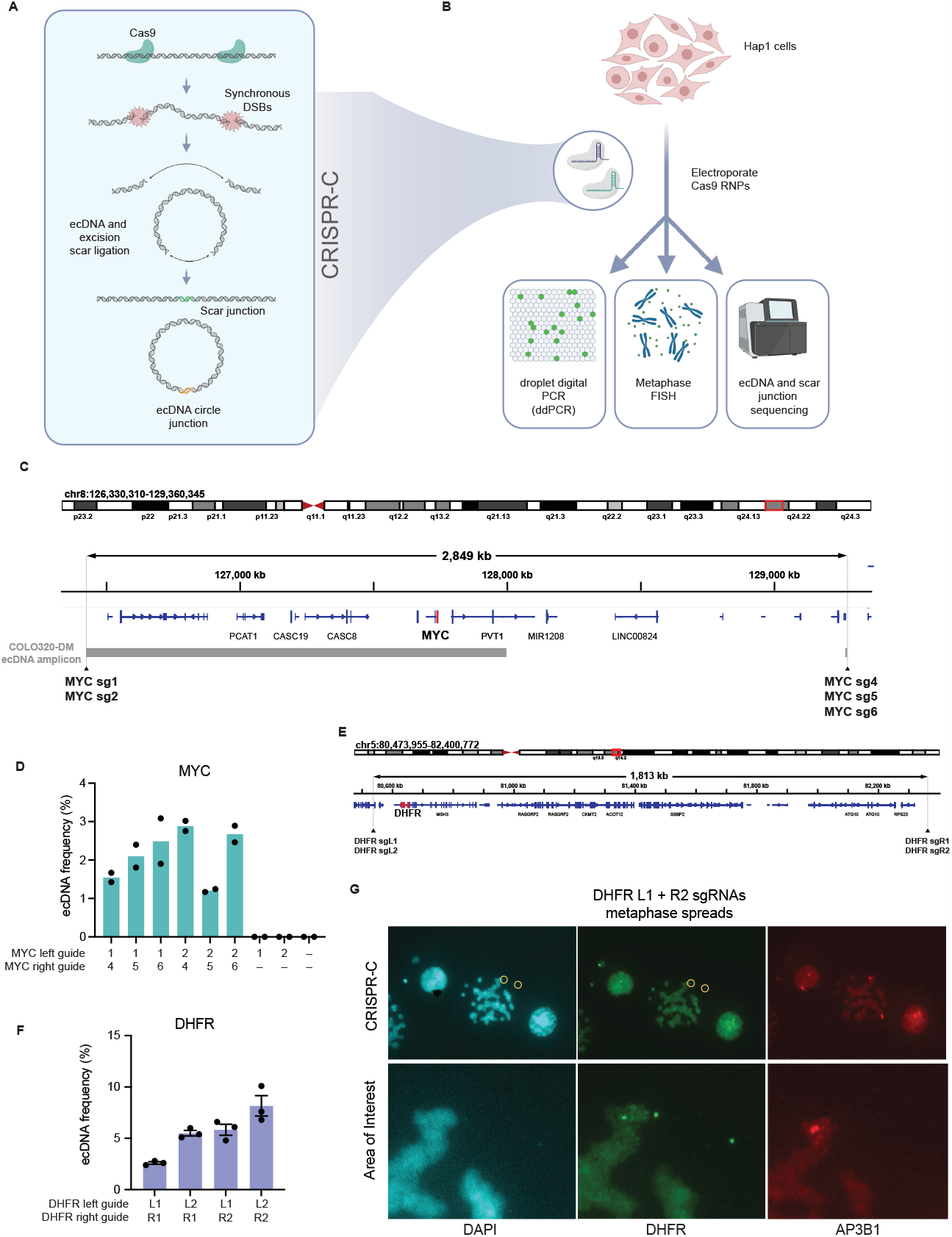
CRISPR-C can generate ecDNA at multiple chromosomal loci. **A**, Schematic depiction of ecDNA generation by CRISPR-C. **B**, Workflow for generating and analyzing induced ecDNA. **C**, Map of chromosome 8 locus indicating positions of guide RNAs used for CRISPR-C, the primary ecDNA amplicon from COLO320-DM, MYC, and other genes within the amplicon. **D**, circularization frequency of different CRISPR-C guide pairs generating MYC-containing ecDNA, as determined by ddPCR. Bars depict means (n = 2 biological replicates). **E**, Map of chromosome 5 locus indicating positions of guide RNAs used for CRISPR-C generating DHFR-containing ecDNA. **F**, circularization frequency of different CRISPR-C guide pairs generating MYC-containing, as determined by ddPCR. Bars depict means, error bars = s.e.m (n = 3 biological replicates). **G**, Metaphase spread fluorescence microscopy images depicting extrachromosomal location of DHFR locus 24-hours post-CRISPR-C. AP3B1 is a DHFR proximal gene on chromosome 5 prior to CRISPR-C.

### 3D chromosome conformation has limited impact on circularization frequency

Translocations occur more frequently between spatially proximal loci, being particularly enriched between intrachromosomal sites(14–16). In *cis*, translocation frequencies correlate with Hi-C contact probability and decrease with linear distance out to approximately 60 Mb(14–17). Single-segment ecDNA generation is a form of intrachromosomal (cis) translocation, and it may be necessary to consider genome conformation when designing CRISPR-C guide RNAs. The end-swapping model predicts that ecDNA formation should be more efficient for spatially proximal loci, as only limited motion is required to merge the repair foci. Conversely, ecDNA formation would be expected to be less influenced by spatial proximity under the dissociation model, as diffusion of legitimate DSB ends away from one another is a defining feature of the model.

EcDNAs range in size from approximately 100 kb to 5 Mb(2), so we focused on the impact of chromosome conformation and Hi-C contact probability on ecDNA generation efficiency between sites separated by ∼1 Mb. We first mapped intrachromosomal interactions in Hap1 cells with Hi-C, identifying an approximately 1Mb loop on chromosome 1 that lacked the conformational complexity of the MYC and DHFR loci (**Fig 2A-C, Supp Fig S2A-B**). We designed guides targeting the ends of the loop, which exhibit high contact frequency between each other, or approximately 1 Mb outside of the loop, which exhibit minimal contact frequencies with the other guide target sites. We performed CRISPR-C using different combinations of these guides, revealing efficient ecDNA generation for all guide pairs tested. Circularization occurred at modestly higher frequencies for guides targeting the sites with high contact probabilities (**Supp Fig 2C**). After normalizing for the activity of the individual guides, however, we found no significant increase in circularization frequency using guides targeting sites with high contact frequencies relative to guides targeting sites with low contact frequencies (**Fig 2D, Supp Fig S2D**). These data suggest that the 3D conformation of the native chromatin has limited impact on the formation of circular ecDNA using CRISPR-C, providing indirect evidence to support the dissociation model.

**Figure 2.**
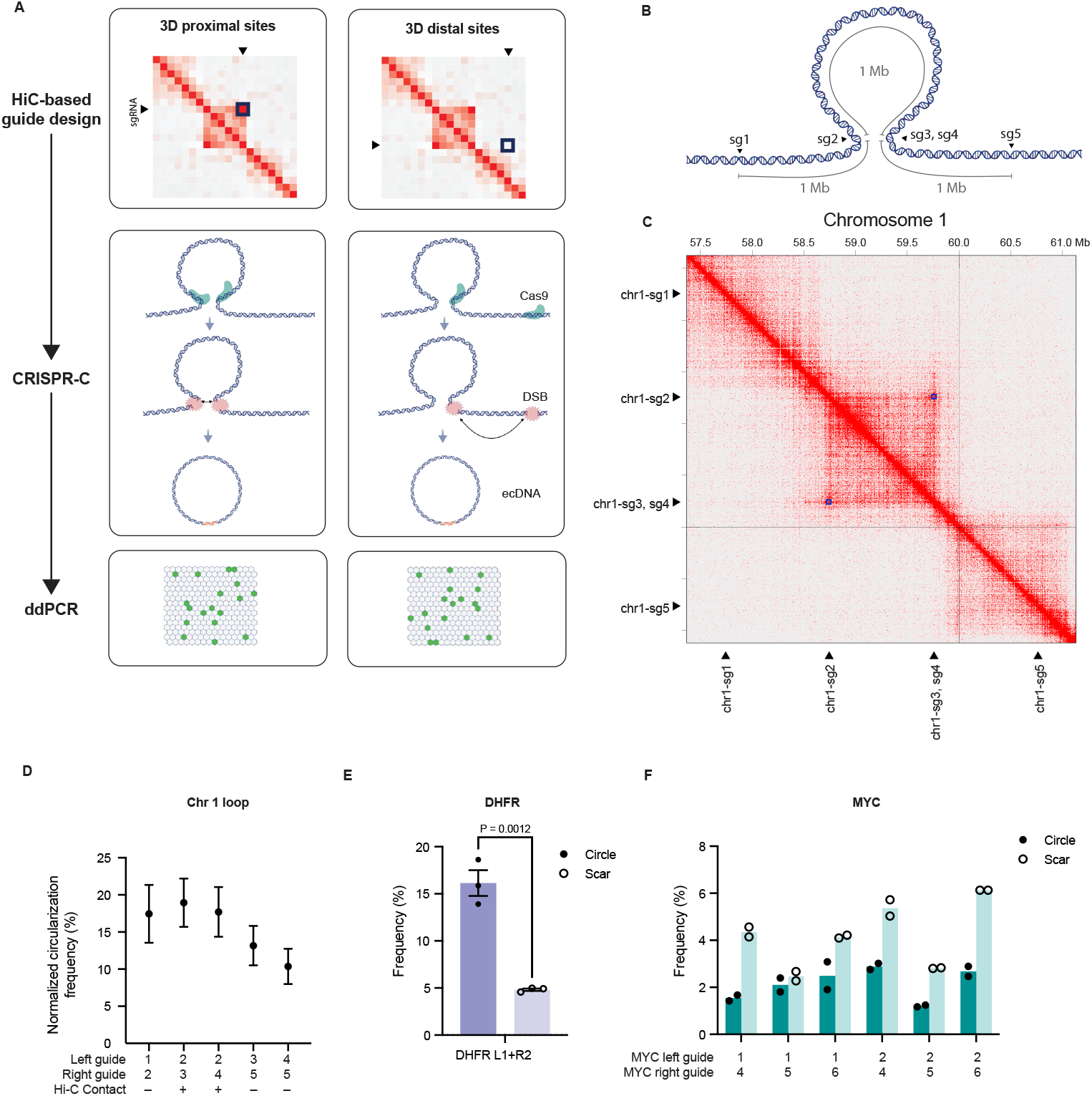
The impact of 3D chromosome conformation on ecDNA formation. Schematic depictions of, **A**, the workflow for examining impact of chromosome conformation on circularization efficiency, and, **B**, the chromosome 1 loop, with the approximate positions of the CRISPR-C guide RNAs used. **C**, Hi-C interaction map of an approximately 1 Mb loop on chromosome 1. Loop indicated by blue squares. Guide locations indicated by black triangles. **D**, normalized circularization efficiency for indicated guide pairs (see methods for calculation), error bars = s.e.m, n = 3 biological replicates. **E**, DHFR ecDNA and scar junction frequencies at 24 hours, reproduced from REF (6), P value calculated with two-sided t-test, n = 3 biological replicates. **F**, ecDNA and scar junction frequencies at 24 hours for MYC CRISPR-C guide pairs in Fig 1C. Bars = mean, n = 2 biological replicates.

### ecDNA and excision scar formation exhibit divergent, site-dependent formation rates

In some ecDNA+ patient-derived samples and cell lines, a deletion exists in the corresponding native chromosomal locus, which we term the “excision scar” (**Fig 1A**)(5,6,18). While the end-swapping model predicts that the excised circle and the excision scar left behind should form at a 1:1 ratio, we previously observed that excision scar formation was less efficient than circularization when generating DHFR ecDNA (**Fig 2E**)(6). To determine if this is a general phenomenon, we quantified the efficiency of excision scar formation for the 6 guide pairs used to generate MYC ecDNA (**Fig 1C, Fig 2F**). As with the DHFR locus, the circle and scar exhibited unequal formation frequencies. However, for the 6 MYC-ecDNA generating guide pairs, scar formation was more efficient than ecDNA formation. These results indicate ecDNA and scar formation are to some degree independent processes, consistent with the dissociation model of ecDNA formation.

### ecDNA and excision scar formation involve NHEJ and MMEJ

While there is evidence of ecDNAs forming via several avenues(7), the mechanisms responsible for ligation to form a covalently closed circle remain unclear. NHEJ is capable of ligating free DSB ends to generate translocations, but its rapidity also opposes translocations(19). Microhomology mediated end joining (MMEJ) can also mediate translocations(20), and microhomologies of 1-5 bp have been found in the ecDNA circle junctions of gliomas (21,22). To examine the contributions of these pathways to ecDNA generation, we performed targeted deep sequencing of ecDNA circle junctions and the excision scar junctions left on the linear chromosome after CRISPR-C (**Fig 3A**).

**Figure 3.**
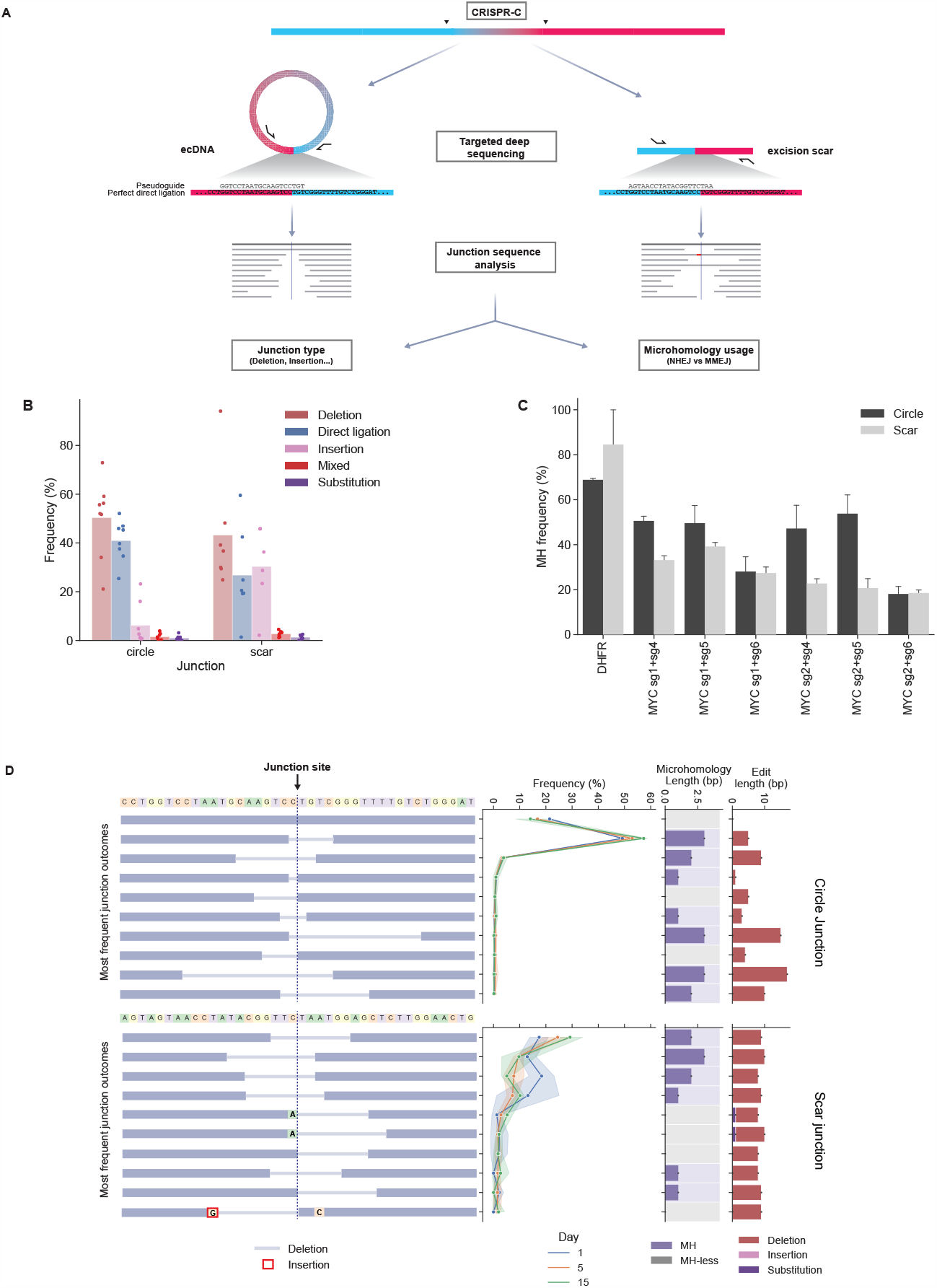
Analysis of junction sequences reveals roles for NHEJ and MMEJ in ecDNA and excision scar formation. **A**, schematic of junction sequencing strategy. **B**, frequency of different junction types for circle and scar junctions. Dots represent mean frequency for each guide pair tested, bars depict mean of all guide pairs. Mixed junctions contain more than one alteration (deletion, insertion, and/or substitution). **C**, Microhomology frequency in circle and scar junctions. Error bars = s.d, n = 3 biological replicates (DHFR and MYC sg2+sg4) and n = 2 for remaining guide pairs. **D**, Frequencies of the 10 most common circle and scar alleles for the DHFR neutral selection timecourse previously reported (6) on indicated days. Error bands = s.d., n = 3 biological replicates. Microhomology length and the length of the indel or substitution for each junction allele are also indicated. Dashed line in left panels indicates junction site.

Interestingly, we found that ecDNA circle junctions and the chromosomal excision scar junctions exhibited distinct ligation sequence characteristics (**Fig 3B**). Although deletions were prevalent in both types of junctions, ecDNA circle junction sequences also exhibited frequent perfect direct ligations, where the two DSBs were ligated without any additional or missing bases, while insertions were rare. By contrast, scar junctions were more likely to contain insertions, while perfect direct ligations were comparatively less common. These different ligation sequence characteristics of the circular ecDNA and the linear chromosome scars suggested that distinct DNA repair processes may underlie the respective formation of their junctions, supporting the dissociation model of excisional ecDNA formation.

To examine the role of microhomology in circle and scar ligation, we classified ligation outcomes as either containing microhomology (MH)—possessing ≥ 1bp of microhomology—or microhomology-less (MH-less). For the purposes of this analysis, all insertions were classified as MH-less. For MH alleles, we also determined the length of microhomology. We found that circle and scar ligation products (junctions) frequently exhibited microhomology, and the frequency of microhomology was highly guide dependent for both circles and scars (**Fig 3C**). Microhomology usage was slightly more prevalent among circle junctions, but this difference was only statistically significant for two guide pairs.

We next examined the allele distributions for each guide pair tested(23). For the DHFR ecDNA, the most common circle junction was a 5 bp deletion exhibiting 3bp of microhomology, while perfect direct ligation was the second most common junction (**Fig 3D**). These two alleles accounted for 70.61% ± 0.55 (s.e.m, n = 3) of all circle junctions. For all guides tested, both circle and scar junctions were dominated by only a few alleles. Generally, the majority of circle junctions comprise the direct ligation product as well as 1-5 MMEJ mediated deletions, except for MYC CRISPR-C pairs involving one specific CRISPR guide (sgRNA6), for which 1bp insertions were common (**Supp Fig S3-4**). The DHFR circle junction sequence distributions tended to remain stable over time under neutral selection pressure, while the scar junction showed more variability across timepoints (**Fig 3D, Supp Fig S5**).

### DNA-PK inhibition promotes ecDNA generation but not excision scar formation

We next sought to investigate whether perturbing DSB repair processes could impact circularization efficiency, as well as further distinguish between the end-swapping and dissociation models of excisional ecDNA formation. In human cells, most DSBs are rapidly repaired by NHEJ, and rapid synapse formation suppresses translocations(19), whereas DSBs that persist have more time to pair with other DSBs and generate translocations(20). The DNA-PK catalytic subunit (DNA-PKcs) is rapidly recruited to DSBs by Ku, which leads to its activation and promotion of NHEJ. While DNA-PKcs phosphorylates many substrates, it’s autophosphorylation appears to be essential for short-range synapsis and NHEJ(19,24). Pharmacologic inhibition of DNA-PKcs inhibits its autophosphorylation, leads to its retention on DSB ends, increases DSB persistence, and ultimately results in an increase in the frequency of translocations (12,20,25–28). We reasoned that treatment with a DNA-PKcs inhibitor would similarly favor CRISPR-C mediated ecDNA generation, which is a form of intrachromosomal translocation. To test this hypothesis, we treated Hap1 cells with 1 uM of the DNA-PKcs inhibitor AZD7648 following CRISPR-C and found that DNA-PKcs inhibition significantly increased ecDNA generation frequency by 2.2-fold (**Fig S6**, 15.23 ± 0.73% vs. 6.65 ± 0.47% in DMSO control, s.e.m., n = 3). Pre-incubation of cells with drug for 1 hour prior to electroporation did not further increase circularization efficiency.

Next, we examined whether inhibition of other DNA damage response (DDR) proteins would affect circularization efficiency. We confirmed that inhibition of DNA-PKcs promoted the formation of ecDNA (**Fig 4A**). Of the other inhibitors tested, only treatment with the ATM inhibitor KU-55933 had a significant effect. ATM inhibition decreased ecDNA formation by 2.5-fold (Figure 3E, 4.40 ± 0.24% vs. 11.02 ± 0.51% s.e.m, n = 3, P = 3.0x10^−4^), whereas inhibition of LIG4, ATR, and PARP (by SCR7, Berzosertib, and Talazoparib, respectively), had no appreciable impact on circularization efficiency.

**Figure 4.**
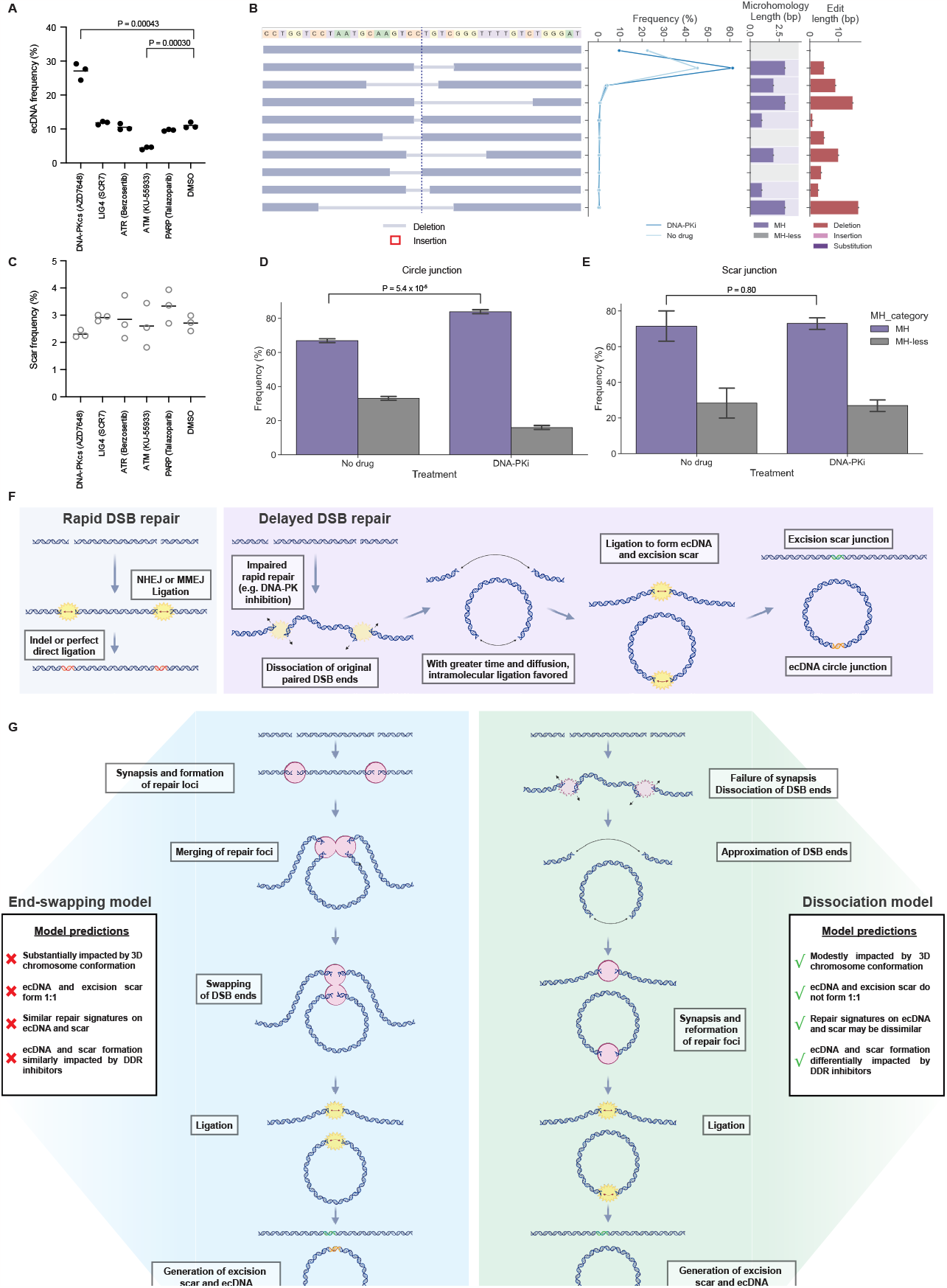
Pharmacologic modulation of ecDNA circularization. **A**, impact of inhibitors of various DNA damage response proteins on CRISPR-C mediated ecDNA generation frequency. Bar indicates mean, n = 3 biological replicates. **B**, the impact of DNA-PKcs inhibition on the distribution of the top 10 DHFR circle junctions. Error bands = s.d., n = 3 biological replicates. **C**, Impact of inhibitors of various DNA damage response proteins on CRISPR-C mediated excision scar frequency. Bar indicates mean, n = 3 biological replicates. Impact of DNA-PK inhibition on, **D**, ecDNA and, **E**, excision scar microhomology usage. Error bars = s.d., n = 3 cell culture replicates. **F**, Schematic of how the rate of DSB repair can promote or disfavor ecDNA formation. **G**, Schematic of two models for excisional ecDNA and scar formation.

In addition to promoting translocations, DNA-PKcs inhibition has been shown to favor MMEJ-mediated repair of CRISPR-induced DSBs over NHEJ(29). Examination of the DHFR ecDNA circle junction sequences demonstrated that DNA-PKcs inhibition decreased the frequency of perfect direct ligation, with a corresponding increase in the frequency of the 5bp MMEJ-mediated junction (**Fig 4B-D, Fig S6**). This shift in pathway usage further validates the involvement of both NHEJ and MMEJ in ecDNA formation, and the ability of these two pathways to compensate for each other in this process. Intriguingly, none of the compounds tested appreciably impacted scar formation frequency (**Fig 4C**). DNA-PK inhibition resulted in a modest decrease in scar formation relative to DMSO, but this effect did not rise to statistical significance (P = 0.086 two-sided t-test). As further evidence of its differential impact on ecDNA vs excision scar formation, DNA-PK inhibition also had little impact on the scar junction sequences or MH usage, in contrast to its promotion of MMEJ at the circle junction (**Fig 4D-E, Fig S6**). The differential impact of both DNA-PK and ATM inhibition on ecDNA and excision scar formation demonstrates they are separate processes, in agreement with the dissociation model.

## DISCUSSION

In this report, we have harnessed CRISPR-C to examine the formation of ecDNAs in human cells, demonstrating that excisional ecDNA formation occurs via the dissociation model rather than the end-swapping model (**Fig 4F-G**). The end-swapping model predicts that (1) circularization of spatially proximal DSBs will be more efficient than between sites with low Hi-C contact frequencies, (2) ecDNA and excision scar formation rates will be near-equivalent, (3) the repair signatures at circle junctions will be similar, and (4) that the impact of DDR inhibitors on ecDNA and scar formation would be similar. All four tests proved contrary to these predictions, which positively supports the dissociation model.

It is well established that spatial proximity, and Hi-C contact frequency specifically, are associated with higher translocation rates(14–17). However, we did not detect significant association of greater Hi-C contact probability with CRISPR-C efficiency, at least for circularizing a 1 Mb chromosomal segment. Previous studies have shown that greater than 90% of intrachromosomal rearrangement breakpoints in breast cancer were within 2 Mb of each other(30), with genome-wide profiling of translocations showing a similar preference for proximal end-joining(16). Thus, it appears that within the typical ecDNA size range (0.1-5 Mb), circularization is sufficiently efficient to not be markedly impacted by variations in 3D conformation. Further characterization in other cell types and across more regions of the genome are needed, but these results indicate that CRISPR-C can be used to efficiently induce ecDNA without first conducting expensive and laborious 3D genome conformation profiling. Moreover, there are fewer constraints on which ecDNA breakpoints can be queried.

In contrast to spatial proximity, we found that manipulation of the temporal dynamics of DSBs could have a profound effect on ecDNA circularization. Small molecule inhibition of DNA-PKcs—known to increase DSB persistence by decreasing DNA-PKcs turnover on DSB ends—led to a dramatic increase in ecDNA generation rates, consistent with the observation that DNA-PKcs promotes translocations between chromosomes(12). Increased DSB persistence likely promotes ecDNA formation by (i) increasing the temporal overlap of the DSBs; (ii) enabling dissociation of the DSB ends from the their legitimate partners; and (iii) with increasing time, favoring the intramolecular ligation event (**Fig 4F**). Recent work has suggested that DNA-PKcs inhibition can decrease formation of ecDNA by chromothripsis, or decrease the genomic span of these ecDNAs (9,31). These seemingly diametric results may be a consequence of a greater reliance on NHEJ to ligate fragments generated by chromothripsis (32). Alternatively, in the context of chromothripsis, where typically several DNA segments are ligated to form an ecDNA, DNA-PKcs inhibition may favor single or oligo-segment circularization events. These smaller ecDNA may be less likely to provide a fitness advantage and are thus neither maintained nor amplified in the cell population.

The DNA damage response has been implicated in ecDNA generation for decades(33), but the processes responsible for ligation of breakpoints to form the circle have not been defined. Both NHEJ and MMEJ have been implicated(21,22,34,35), however their relative contributions have been unclear. Our work has revealed that both NHEJ and MMEJ are capable of performing the critical repair and ligation steps of ecDNA generation. The proficiency of each pathway to play this role suggests that the ligation step is robust, with either pathway able to compensate for inhibition or deficiency of the other. Thus, this may be a difficult step to target therapeutically. Indeed, we found that pharmacologic manipulation of several DDR factors had no appreciable impact on ecDNA generation. The lone exception, besides DNA-PKcs, was ATM inhibition, which decreased ecDNA generation efficiency. ATM has been termed the “master regulator” of the DSB response(24), phosphorylating hundreds of substrates in the process of orchestrating orchestrating the cellular response to DSBs. ATM inhibition with KU-55933 has been shown to limit DSB mobility(36) and abrogate DSB clustering(37). Both of these effects of ATM inhibition would be expected to antagonize ecDNA formation (**Fig 4F-G**).

Contrary to expectations(7), our results demonstrate that the circularization of a chromosomal segment between contemporaneous DSBs can be remarkably efficient, reaching 15% in some cases, or even higher in the context of DNA-PKcs inhibition. Beyond the implications for the use of CRISPR-C as a research tool, this observation suggests that excisional ecDNA formation is likely far more common than previously appreciated(7), and that circularization of contemporaneous DSBs is likely not the rate limiting step in ecDNA formation. Further work is needed to determine the frequency of contemporaneous DSBs forming through endogenous processes or environmental insults (radiation, chemotherapeutics) within 0.1-5 Mb.

In this study, we have demonstrated the power of ecDNA generation with CRISPR-C to yield meaningful insights into ecDNA biogenesis, establishing the dissociation model of excisional ecDNA formation, and revealing mechanistic divergence in the formation of ecDNA and their chromosomal scars. Future therapeutic strategies to forestall the emergence of ecDNAs may target not only the mechanism of ecDNA formation, but the distinct processes governing repair of the linear locus from which they were excised. Here, we used the near-haploid Hap1 cell line, which is an established tool for genetic screens and other studies(38–40), but it will be necessary in the future to extend our work to a range of non-haploid cell-types and more genomic sites. Aided by the portability of CRISPR-C, we anticipate this approach will permit distinct and powerful investigations of the entire lifecycle of ecDNAs in various contexts.

## METHODS

### Cell culture

Hap1 cells (Horizon Discovery) were maintained in IMDM supplemented with GlutaMAX and 10% FCS (Gibco). Cells regularly tested negative for mycoplasma contamination.

### Guide RNA design for CRISPR-C

sgRNAs were designed to target regions corresponding to the span of the canonical MYC-containing ecDNA from the COLO320-DM cell line as in Fig 1B. Guides for generating DHFR-containing ecDNA were previously described(6). To examine the role of chromosome conformation on CRISPR-C, regions of interest were identified by examining the HiC contact matrix (Fig 2B). For all guides, regions of interest were examined within the UCSC genome browser and RepeatMasker and GeneHancer tracks were used to avoid targeting repetitive regions and regulatory elements, respectively. Guides were designed using the Integrated DNA Technologies (IDT) Custom Alt-R CRISPR–Cas9 guide RNA software (https://www.idtdna.com/site/order/designtool/index/CRISPR_CUSTOM). These sequences were ordered as Alt-R sgRNAs (IDT).

### ecDNA induction by CRISPR-C

Hap1 cells were trypsinized, quenched with IMDM (GlutaMAX, 10% FCS), counted and centrifuged at 300g for 5 min. Cells were washed once with PBS before resuspension in Neon Resuspension Buffer to 1.1 × 107 cells ml−1. Ribonucleoprotein (RNP) complexes were formed as follows: Cas9 (IDT) was diluted to 36 μM in Neon Resuspension Buffer. Equal volumes of diluted Cas9 and sgRNA (44 μM in TE, pH 8.0) were mixed and incubated at room temperature for 10–20 min. Left and right sgRNA RNPs were assembled separately. Then, 5.5 μl of each RNP, 5.5 22 μl of electroporation enhancer (10.8 μM; IDT) and 99 μl of cells were mixed and electroporated according to the manufacturer’s instructions using a 100-μl Neon pipet tip and electroporated with the Neon Transfection System (Thermo Fisher Scientific) using the following settings: 1,575 V, 10-ms pulse width, 3 pulses. Single-guide controls were prepared as above except 11 μl of the appropriate sgRNA was used. Volumes were scaled down for 10 uL electroporations. Cells were collected at the 24 hours post-electroporation as follows: cells were washed with 1 ml per well prewarmed PBS (Gibco), followed by the addition of 100 μl TrypLE Express (Thermo Fisher Scientific) and incubation at 37 °C for 5–10 min. TrypLE was quenched with 800 μl IMDM (GlutaMAX, 10% FCS) and the cell suspension was pelleted at 300 g for 5 min at 4 °C. The supernatant was discarded and the cell pellets were stored at −80 °C.

For experiments including inhibition of DDR proteins, after electroporation, cells were plated in media containing either AZD7648 (1 μM), SCR7 (1 μM,), Berzosertib (80 nM), KU-55933 (10 μM) Talazoparib (10 nM), or DMSO. Samples were harvested 24 hours after electroporation. All compounds were obtained from Selleck Chemicals. Final DMSO concentration in all wells was 0.2%. Final drug concentrations were selected based on effective doses for cell culture reported in the literature(41–45).

### ddPCR to determine ecDNA or chromosomal scar frequency

gDNA was isolated using DNeasy columns (QIAGEN) according to the manufacturer’s instructions, including a 10-min incubation at 56 °C during the proteinase K digestion step; DNA was eluted with 100 μl EB buffer. Amplicons for the ecDNA junction, chromosomal scar junction and glyceraldehyde 3-phosphate dehydrogenase (GAPDH) were designed using the IDT PrimerQuest software (https://www.idtdna.com/PrimerQuest/Home/Index). Dual-quenched probes (IDT) were used: FAM-labeled probes were used for both the ecDNA and chromosomal scar junction amplicons to facilitate multiplexing with the GAPDH amplicon utilizing a HEX-labeled probe. All probe and primer sequences are available in Supplementary Data Set 1. Droplets were created using droplet-generating oil for probes, DG8 cartridges, DG8 gaskets and the QX200 Droplet generator (Bio-Rad Laboratories); amplification was performed using the ddPCR Supermix for Probes (Bio-Rad Laboratories). The ddPCR Supermix amplification reactions were set up according to the manufacturer’s specifications (Bio-Rad Laboratories). Approximately 60 ng of gDNA was used in a 20 μl reaction with a final primer concentration of 900 nM (225 nM for each primer), 125 nM FAM probe and 125 nM HEX probe. The reaction was partitioned into droplets for amplification according to the manufacturer’s protocol (Bio-Rad Laboratories). Droplets were transferred to a 96-well PCR plate and heat-sealed using the PX1 PCR plate sealer (Bio-Rad Laboratories). Droplets were amplified using the following cycling conditions: 95 °C for 10 min, 40 cycles (94 °C for 30 s, 56.1 °C for 60 s), 98 °C for 10 min. After thermal cycling, droplets were scanned individually using the QX200 Droplet Digital PCR system (Bio-Rad Laboratories). Positive and negative droplets in each fluorescent channel (HEX, FAM) were distinguished on the basis of fluorescence amplitude using a global threshold set by the minimal intrinsic fluorescence signal resulting from imperfect quenching of the fluorogenic probes (negative droplets) compared to the strong fluorescence signal from cleaved probes in droplets with amplified template(s). The frequency of ecDNA or chromosomal scar was calculated by dividing their measured concentration by the concentration of the GAPDH amplicon. Normalized circularization frequency (Fig. 2D) was calculated as follows:

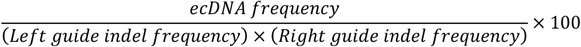

The indel frequencies were determined for each guide individually as below.

### CRISPR editing quantification

Quantification of editing by CRISPR with individual sgRNAs was performed using the ICE analysis tool (Synthego, v3.0). Briefly, the target locus was amplified using Platinum SuperFi polymerase master mix (Thermo Fisher Scientific; for primers see Supplementary Data Set 1). PCR purified and sanger sequencing were performed by Elim Biopharmaceuticals, Inc. Sequence traces and sgRNA spacer sequences were then uploaded to the ICE webtool for analysis.

### Metaphase FISH

Cells were arrested in mitosis with KaryoMAX™ Colcemid™ Solution (Gibco, #15122012) for 4 hours. The cells were then collected and washed once in 1X PBS. The cell pellet was resuspended in 50μL 1X PBS and treated with 0.075M KCl buffer for 20 mins at 37°C. The cells were then fixed in fresh Carnoy’s fixative (3:1 methanol:glacial acetic acid) followed by three additional washes with the fixative. Cells were then dropped to a humidified slide and air-dried at room temperature. The slide was briefly equilibrated in 2X SSC buffer and subjected to dehydration with ascending series of ethanol (70%, 85%, 100%) for 2 mins each. The slides were air dried completely and FISH probes (Empire Genomics, DHFR gene: DHFR-20-GR, AP3B1 gene: AP3B1-20-RE) diluted in 1:6 ratio with hybridization buffer were applied to the sample. A coverslip was applied and the sample was subjected to heat denaturation at 75°C for 3 mins, followed by hybridization overnight at 37°C. The coverslip was removed and the slide was washed in 0.4X SSC buffer and then in 2X SSC (0.1% Tween) for 2 mins each. DAPI stain (50ng/mL) was used to stain for nuclei for 2 mins and the slides were washed once briefly in ddH_2_O. The slides were mounted with Prolong Diamond Antifade (Invitrogen, #P36961) and air dried overnight. Images were acquired on a Leica DMi8 Thunder imager using a 63X oil objectives.

### Targeted deep sequencing of ecDNA and excision scar junctions

20 cycles of primary PCR to amplify the region of interest was performed using ∼200ng μL of genomic DNA in a 10 μL Platinum SuperFi II polymerase reaction (Thermo Fisher Scientific; for primers see Supplementary Data Set 1). Illumina adapters and indexing sequences were added via 15 cycles of secondary PCR with 1 μL of primary PCR product in a 10 μL Platinum SuperFi II polymerase reaction. The final amplicons were run on a TAE-agarose gel (0.7%); and the product band was excised and extracted using the Freeze and Squeeze Kit according to the manufacturer’s instructions (Bio-Rad). Gel-purified amplicons were quantified with a dsDNA HS Assay kit on a Qubit fluorometer (ThermoFisher Scientific). Then, amplicons were pooled and sequenced on the NextSeq 550 platform (Nextseq 500/550 Mid Output Kit v2.5, Illumina, for primers see Supplementary Data Set 1).

### Sequencing analysis

Analysis of junction sequences was performed with CRISPResso2(46) in batch mode utilizing paired-end reads with the following command:

CRISPRessoBatch --batch_settings <batch_settings_file> -q 30 -n <output_name> --ignore_substitutions

For each junction, the expected product of perfect direct ligation, in which the Cas9 cutsites of each guide are directly ligated without additional loss or insertion of sequence, was used as the reference sequence, and a “pseudo-guide” sequence was used whose cut site aligned to the junction (Fig 3A).

Length of microhomology was quantified for each junction sequence as follows: the last *n* nucleotides of the upstream sequence were compared to the last *n* nucleotides of the deleted sequence. The largest *n* for which there was an exact sequence match was definied as the upstream microhomology, *mh1*. This process was repeated to compare the first *n* nucleotides of the deletion to the first *n* nucleotides of the downstream process to calculate the downstream microhomology, *mh2*. The microhomology length was identified as the greater of *mh1* vs *mh2*. Junctions with microhomology length ≥ 1 were defined as exhibiting microhomology (MH), while deletions with no microhomology and insertions were defined as microhomology-less (MH-less)(47,48).

### Hi-C

Hi-C was performed by following the reported HiChIP protocol and omitting the protein immunoprecipitation step(49). Briefly, 10^6^ cells were crosslinked with 1% formaldehyde for 10 minutes. Formaldehyde was quenched by the addition of glycine. Nuclei were isolated in lysis buffer (1% Triton x-100, 0.1% SDS, 150 mM NaCl, 1mM EDTA, and 20 mM Tris, pH 8.0) by rotating the cells for 1-hour at 4C. Nuclear pellet was resuspended in 0.5% SDS solution and incubated at 62C for 10 minutes. SDS was quenched by the addition of 10% Triton-X. Chromatin was digested with 8U of MboI enzyme overnight at 37C. Biotin fill-in was performed for 1-hour at 37C, followed by proximity ligation for 6-hours at room temperature. After proximity ligation, nuclei were subjected to Proteinase K digestion and decrosslinking overnight at 68C. DNA was column purified (Minelute, Qiagen), and sonicated to a size distribution of 200-500bp. DNA was subjected to streptavidin bead binding to pull-down biotinylated DNA that represents ligation junctions. Tn5 was used to create libraries by on bead tagmentation with sequencing adapters, followed by PCR. Libraries were quantified by Qubit and size distribution was assessed with Bioanalyzer. Libraries were sequenced on NovaSeq 6000 with paired-end 150bp configuration.

### Hi-C Analysis

Paired end reads were aligned to the reference genome (hg38) using the HiC-Pro pipeline (version 2.11.0). Default settings were used to remove duplicate reads, assign reads to MboI restriction fragments, filter for valid interactions, and generate binned interaction matrices. The Juicer pipeline’s HiCCUPS tool was used to identify loops. Filtered read pairs from the HiC-Pro pipeline were converted into .hic format files and input into HiCCUPS using default settings. Contact matrices were visualized using Juicebox using balanced normalization.

### Statistics and reproducibility

Unpaired two-sided t-tests were performed using the scipy stats package v1.10.1 and Graphpad Prism v9.5.1. Schematics in Figures 1A-B, 2A-B, and 4F-G were created with BioRender.

## Data availability statement

Sequencing data are available at the NCBI Short Read Archive ([*Data to be uploaded prior to publication*]). All other data that support the findings of this study are available from the corresponding authors upon request.

## ACKNOWLEDGEMENT

We thank members of the Chang and Mischel labs for discussion. This project was supported by Cancer Grand Challenges CGCSDF-2021\100007 with support from Cancer Research UK and the National Cancer Institute (H.Y.C., P.S.M.). H.Y.C. is an Investigator of the Howard Hughes Medical Institute. J.C.R. is supported by NIH K99-CA279512 and the A.P. Giannini Foundation.

