## Supplemental Information for "Disparate pathways for extrachromosomal DNA biogenesis and genomic DNA repair"

### Supplementary Info

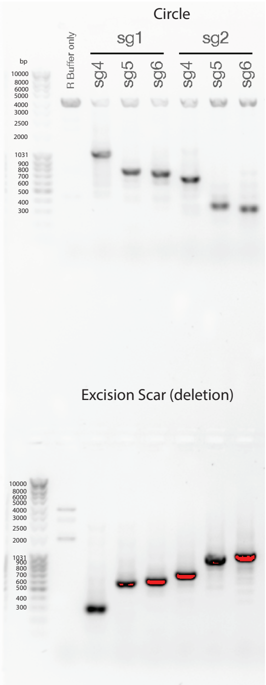


| Expected product size with no substantial deletion or insertion | | | |
|---|---|---|---|
| | | Size (bp) | |
| | | Primers | |
| Left sgRNA | Right sgRNA | CIRCLE | DELETION |
| sg1 | sg4 | 1051 | 309 |
| sg1 | sg5 | 747 | 613 |
| sg1 | sg6 | 714 | 646 |
| sg2 | sg4 | 635 | 725 |
| sg2 | sg5 | 331 | 1029 |
| sg2 | sg6 | 298 | 1062 |

**Figure S1. PCR detects circle junctions for all MYC ecDNA guide pairs tested.** Gel electrophoresis analysis of MYC CRISPR-C guide screen and the expected sizes of circle (ecDNA) and deletion (excision) scar products for all guide pairs tested. R Buffer only cells were resuspended in electroporation buffer but not electroporated.

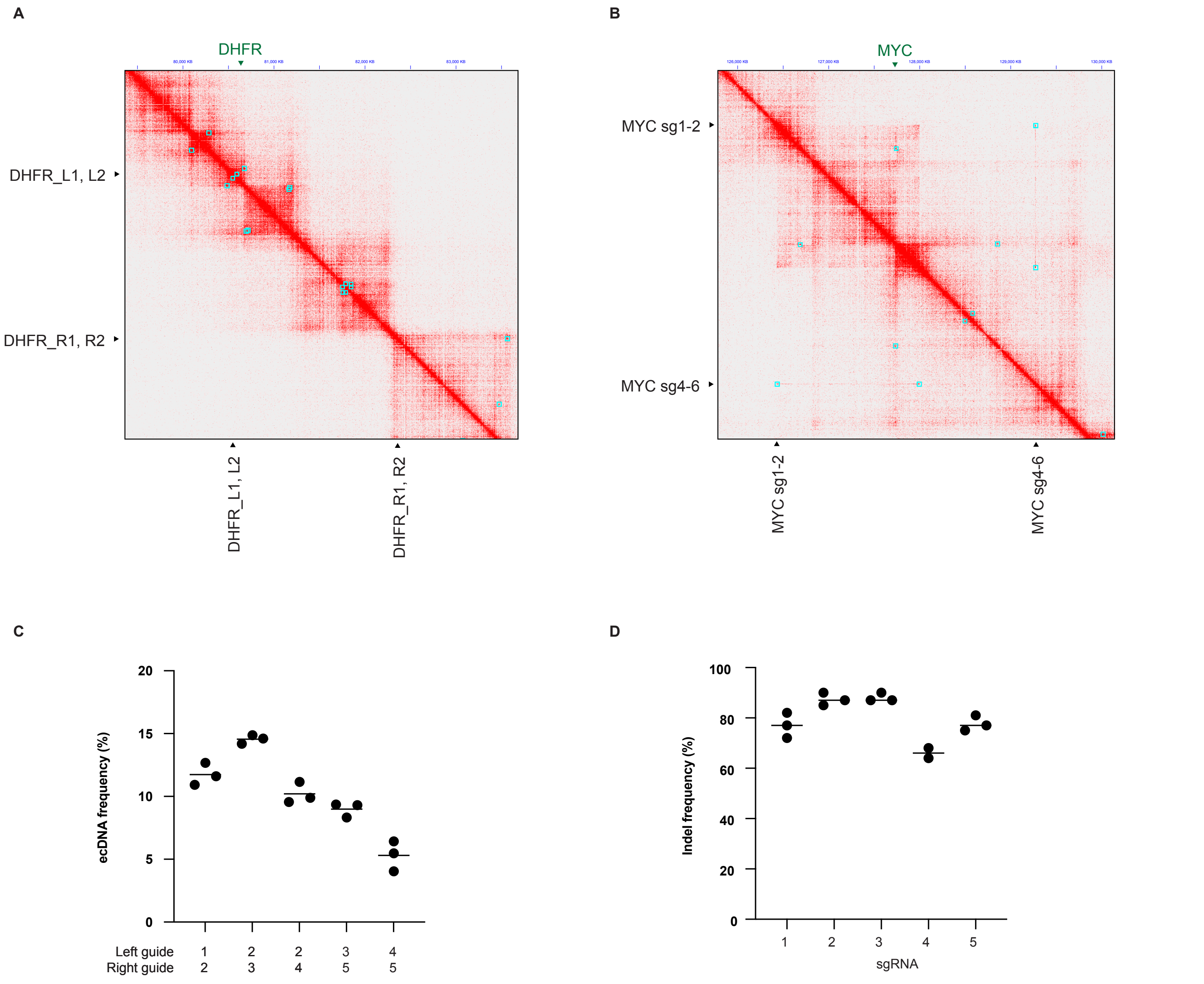


**Figure S2. MYC and DHFR loci Hi-C contact maps & unnormalized Hi-C CRISPR-C data.** Hi-C interaction map of, **A,** DHFR on chromosome 5 and, **B,** MYC loci on chromosome 8. Loops indicated by blue squares. Guide locations indicated by black triangles. **C,** ecDNA junction frequency for indicated guide pairs targeting the chromosome 1 loop loci in Fig. 2 A-C. Line indicates mean, n = 3 biological culture replicates. **D,** editing efficiency of individual guides in C, as determined by sanger sequencing and analysis with the Synthego ICE web tool. Line indicates mean, n = 3 biological culture replicates, except for sgRNA4 (n = 2).

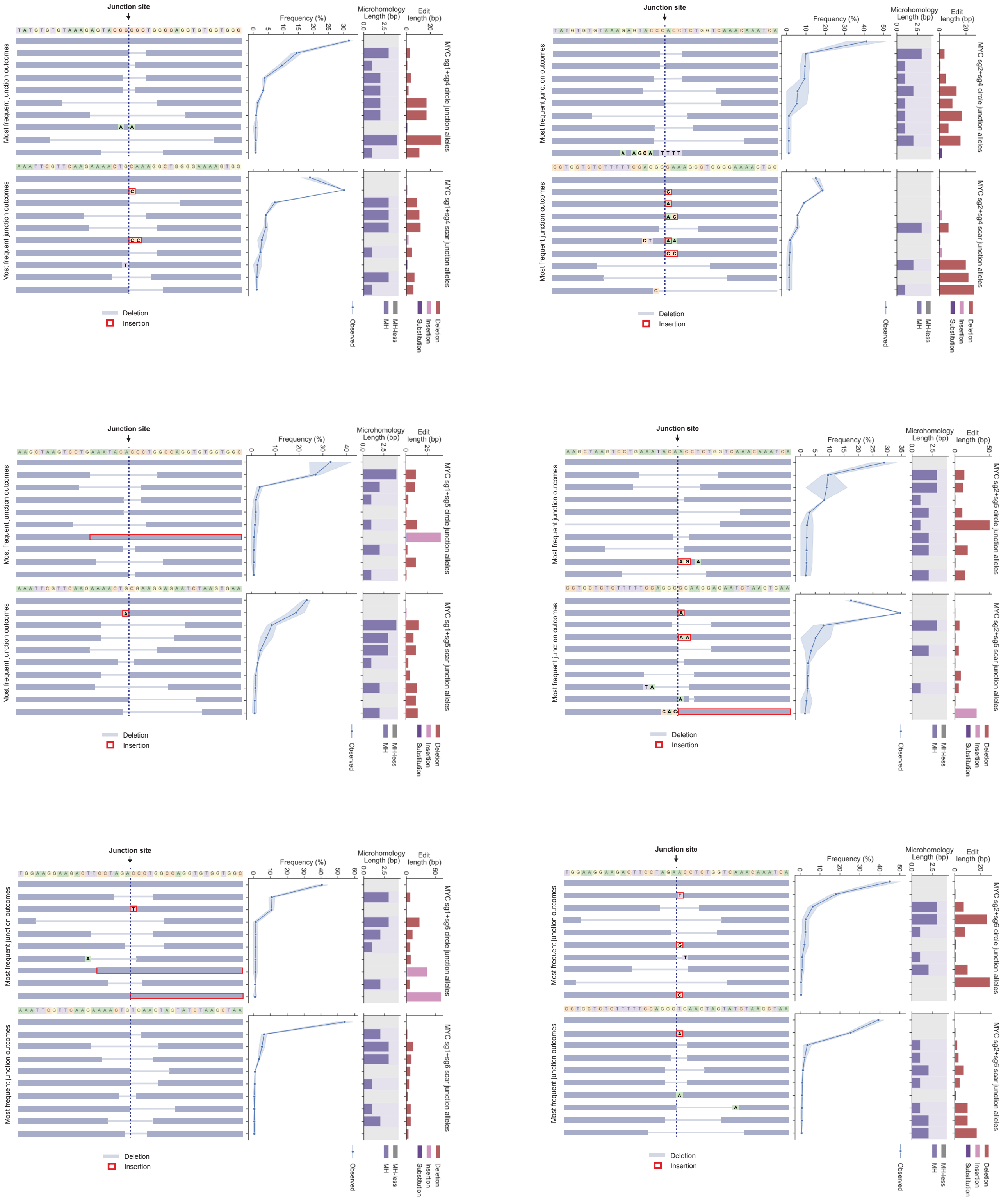


**Figure S3. Junction profiles for MYC CRISPR-C guide screen.** Frequencies of the 10 most common circle and scar alleles for all MYC CRISPR-C guide pairs tested in Fig 1C . Error bands = range, n = 2 biological replicates. Microhomology length and the length of the indel or substitution for each junction allele are also indicated. Dashed line in left panels indicates junction site.

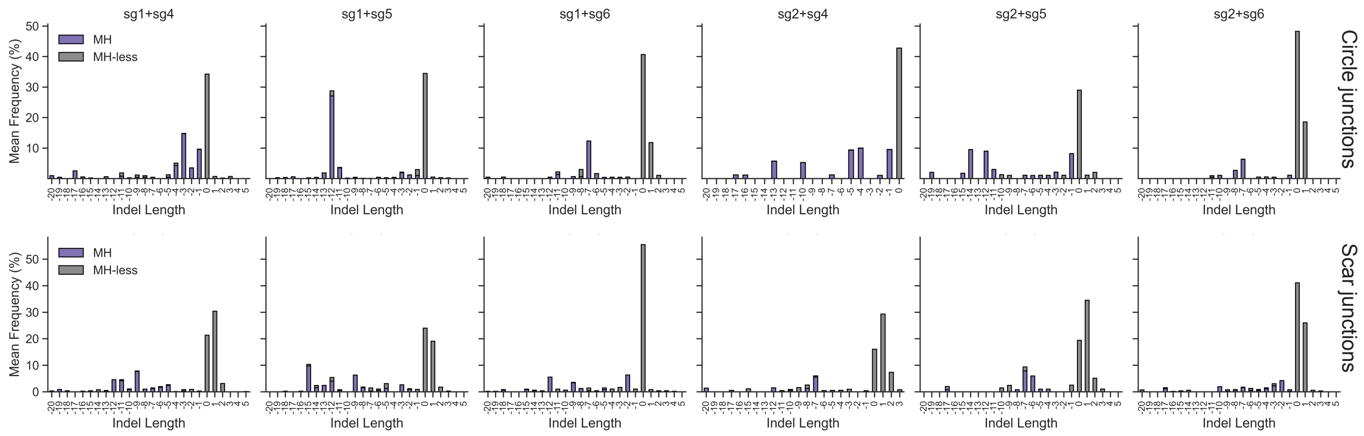


**Figure S4. Junction indel size for MYC CRISPR-C guide screen.** Mean observed and predicted indel size distribution for circle and scar junctions of all MYC CRISPR-C guides tested in Fig 1C.

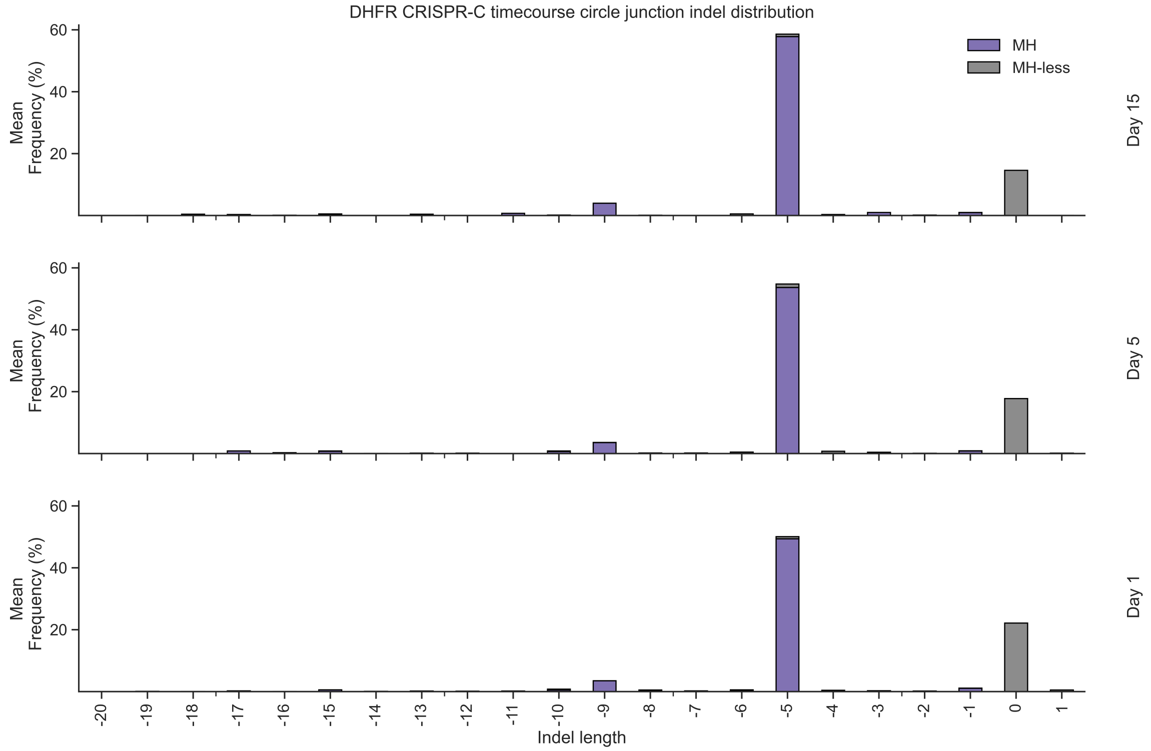


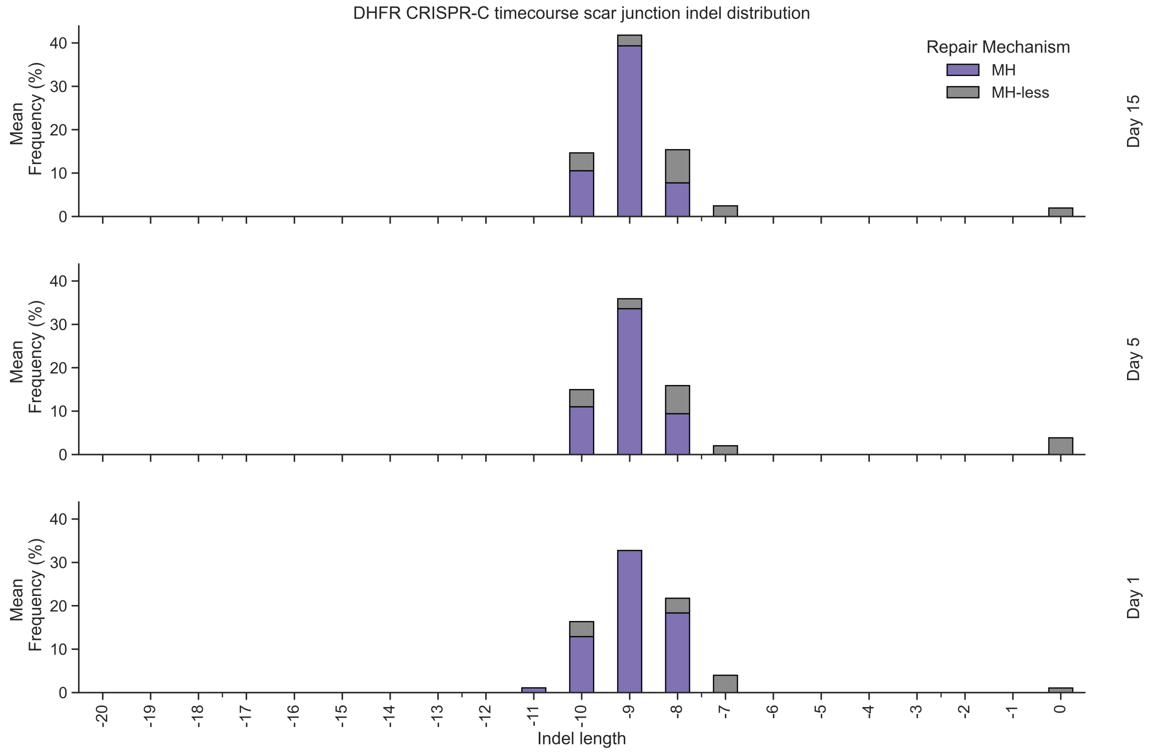


**Figure S5. DHFR timecourse ecDNA and excision scar junction indel size distributions are stable over time.** Mean observed indel size distribution for DHFR circle and scar junction at 24 hours, day 5, and day 15.

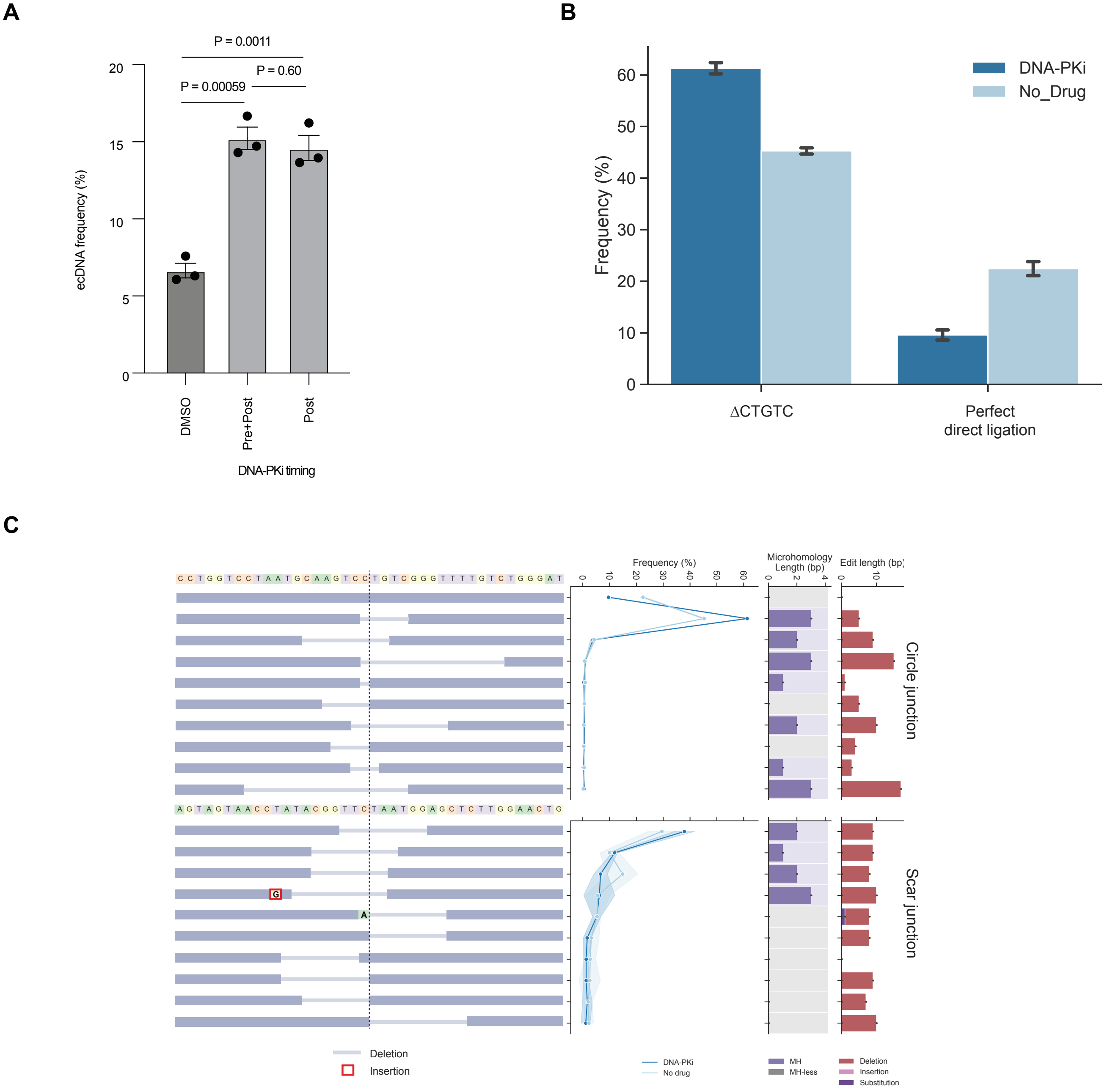


**Figure S6. DNA-PK inhibition promotes ecDNA formation, favors MMEJ in ecDNA formation, but has minimal impact on excision scar junctions. A,** The impact of treatment of Hap1 cells for 1 hr before and 18 hours after CRISPR-C with 1 uM of the DNA-PK inhibitor AZD7648 (Pre+Post), 18 hours of treatment after CRISPR-C with no pre-treatment (Post), or DMSO. Error bars = s.e.m., n = 3 biological replicates. P values calculated using a two-sided t-test. **B,** The impact of DNA-PK inhibition (Post) on the two most common junctions: a 5 bp deletion exhibiting 3 bp of microhomology (ΔCTGTC) and perfect direct ligation. Error bars = s.d, n = 3 cell culture replicates. **C,** the impact of DNA-PKcs inhibition on the distribution of the top 10 DHFR circle and scar junctions. Error bands = standard deviation, n = 3 biological replicates. Microhomology length and the length of the indel or substitution for each junction allele are also indicated. Dashed line in left panels indicates junction site. Circle junction distribution is also displayed in Figure 4B.
